# Modeling Protein Structure Using Geometric Vector Field Networks

**DOI:** 10.1101/2023.05.07.539736

**Authors:** Weian Mao, Muzhi Zhu, Hao Chen, Chunhua Shen

**Affiliations:** Zhejiang University, China; The University of Adelaide, Australia

## Abstract

Proteins serve as the foundation of life. Most diseases and challenges in life sciences are intimately linked to protein structures. In this paper, we propose a novel vector field network (VFN) for modeling protein structure. Unlike previous methods that extract geometric information relying heavily on hand-crafted features, VFN establishes a new geometric representation paradigm through a novel vector field operator. This vector field operator can not only eliminate the reliance on hand-crafted features, but also capture the implicit geometric relationships between residues. Thus, it enables VFN to have better generalizability and flexibility. We evaluate VFN on the protein inverse folding task. Experiment results show that VFN can significantly improve the performance of the state-of-the-art method, PiFold, by 3.0% (51.7% *vs*. 54.7%) in terms of the sequence recovery score, and outperform the recent solid baseline, Protein MPNN, by 8.7% (46.0% *vs*. 54.7%). Furthermore, we scale up VFN with all known protein structure data. Finally, the model achieves a recovery score of **57.1**%, pushing the accuracy to the next level.

## 1 Introduction

Proteins are the fundamental building blocks of life and play crucial roles in biological systems. Many challenges and diseases in the life sciences are intricately related to proteins. Insights into the structures of proteins can help to develop treatments, to understand diseases, and to elucidate the principles of life.

In this paper, we propose a machine learning model for modeling protein structure, termed vector field networks(VFN), which aims to develop a simpler, more generalizable and efficient structure modeling approach. Here, we select the inverse protein folding task [20] as a benchmark to evaluate the model’s performance; because other tasks, *e*.*g*., protein structure prediction, often associate with some confounding factors, *i*.*e*., MSA and hand-designed energy function [39]. As such, the inverse folding task provides a more rigorous evaluation for protein structures model, free from the influences of external information sources. Moreover, the inverse protein folding task is a critical step in the *de novo* protein design process and has important implications for applications such as antibody drug development.

For the inverse protein folding problem, numerous protein 3D structure encoders have been proposed. Recently, the best-performing methods have almost all adopted the paradigm of combining message-passing neural networks [16] (MPNNs) with hand-crafted geometric features, *e*.*g*., the distances between atom pairs within near residues. However, those approaches may suffer from two limitations: 1) *These models lack the ability to extract geometric information. Thus, the expressive flexibility is limited*. Furthermore, those methods have to rely on the manually-designed featurizer, while hand-crafted features are difficult to cover all the geometric features, such as disulfide bonds. This limitation may prevent those methods from achieving better performance. 2) *The generality of hand-crafted features is limited*. Because, in more complex tasks, *e*.*g*., protein structure prediction [12, 23, 29] or drug discovery [50], hand-crafted features are required to encode not only the backbone atoms but also a considerable number of side chain atoms in some residue, *i*.*e*.,26 atoms in the arginine in total. In such cases, very sophisticated features are inevitably needed, which pose significant implementation challenges for featurizers, *i*.*e*.,676 atom pairs existing between two arginines.

To alleviate these limitations, we propose the geometric vector field network. The core idea of VFN is that, when modeling the geometric relationship between two amino acids, an operator is designed to establish a vector field in Euclidean space for both residues. As a result, certain implicit vector representations of the two amino acids can be employed to model the spatial relationship through this vector field operator. *This allows VFN to learn to extract geometric features*. Furthermore, VFN is *more flexible and more generalizable*. Specifically, the introduction of vector fields allows the model to avoid relying on complicated featurizers. Since part of the vector representations of amino acids are initialized as the atomic coordinates of amino acids, the explicit geometric relationships between these atoms can be captured through the vector field operators. Those explicit geometric relationships actually provide the information that was provided by featurizers in the past. Moreover, the geometric information that the featurizer failed to capture may also be modeled by VFN.

Compared with the GVP [22], which also uses vector representations, VFN differs mainly in two aspects: 1) VFN directly embeds the implicit vector representations in the Euclidean space as virtual atoms, refer to the top of Fig.1(B). The geometric constraints between those virtual atoms of two amino acids are strictly guaranteed by our vector field operators. However, in GVP, the vector representations are only the implicit variables in the neural network and do not satisfy the geometric constraints as in VFN. 2) To satisfy the SE(3) equivariance, GVP only uses the operators which satisfy SE(3) equivariance for vector representations, *e*.*g*., *L*_2_ normalization. Some basic operators, *e*.*g*., ReLU [1], are not allowed for vector representation. This may limit the representational capacity of the model. In contrast, VFN transforms all vector representations into the local residue frame to satisfy SE(3) invariance without restrictions on the operators of the model. As a result, various operators, which do not satisfy the SE(3) equivariance restrictions, can be proposed and used in VFN. Those operators improve the representational capacity of VFN, proven by the experiments as shown in Table 3.

**Figure 1:**
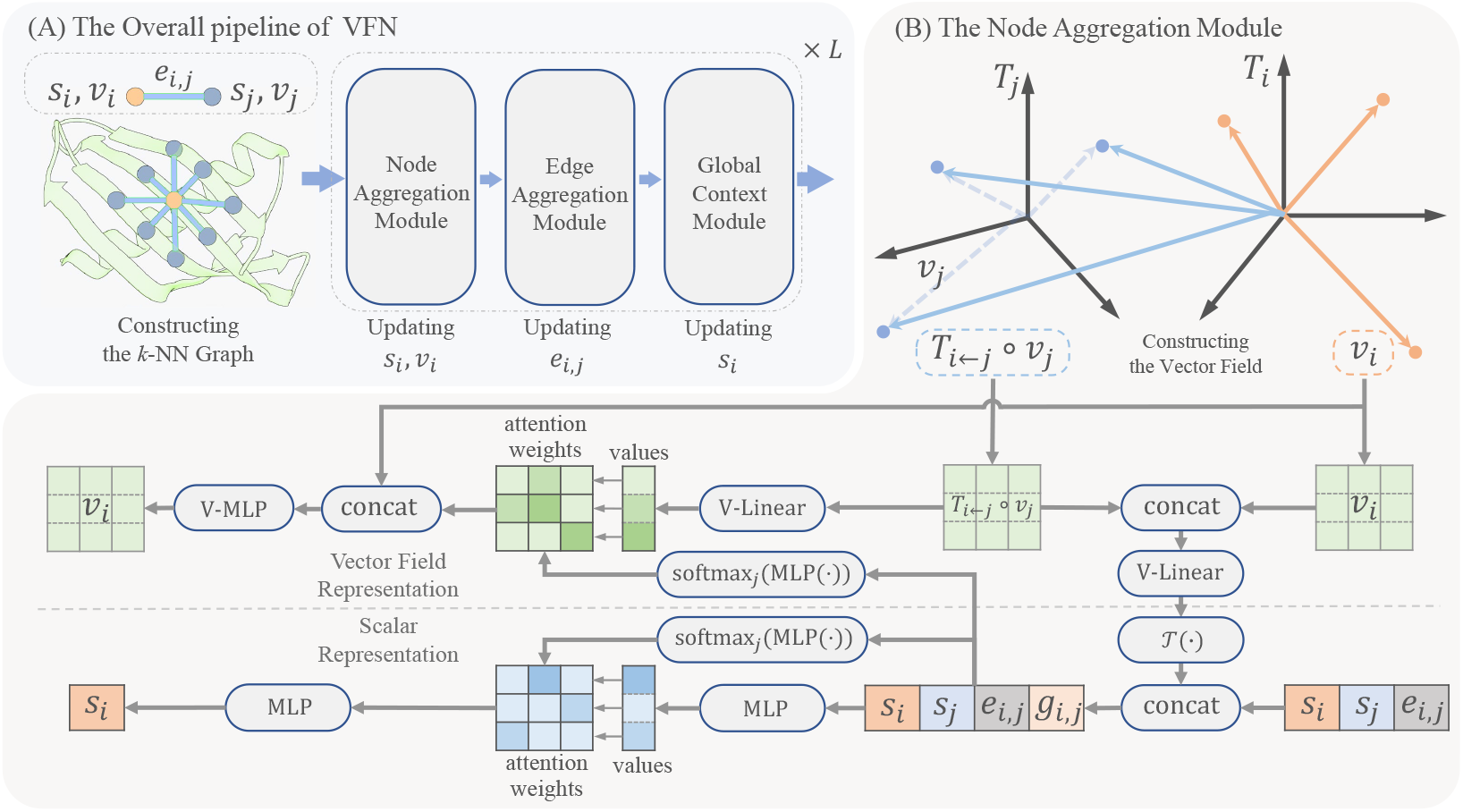
An overview of VFN. Figure (a) shows the overall pipeline of VFN. Figure (b) details the specific functions of the Node Aggregation Module, which serves as the pivotal component of VFN. The specific notations are elaborated in §3.2.

Based on inverse folding, experimental results show that the VFN has achieved the best performance to date on the CATH 4.2 dataset, reaching an accuracy of 54.7% even without the use of edge featurizers. Moreover, compared to protein MPNN, our performance has improved by 8.7%. Furthermore, we have scaled up our model with more data, and achieved an accuracy of 57.1% on the same test set. Our contributions can be summarized as follows:

- We propose VFN, which obviates the reliance on hand-crafted features. The benefits are evident. On the one hand, it eliminates the need for complex hand-crafted feature design for more complicated tasks, making it more versatile. On the other hand, the model can capture implicit geometric features through vector fields.
- VFN has shown significant improvements in inverse folding tasks. It has achieved SotA performance even without edge featurizers, surpassing the current SotA performance by 3.0%. By scaling up with more training data, the model’s accuracy was further improved to 57.1%, providing a more accurate tool for protein design.
- The proposed vector field based on local residue frame representation provides a new paradigm for future protein structure modeling methods.

## 2 Related Work

### 2.1 Representation of Protein Structure

Proteins consist of amino acids, of which there are 20 distinct types found in nature. Amino acids have a conserved backbone composed of four atoms: *C, C*_*α*_, *N, O*, shared by all kinds of amino acids. In previous work, some hand-crafted geometric features are proposed to represent the structure of a protein based on the coordinate of backbone atoms. Those hand-crafted features [20, 14] include the distance and direction between the atoms in different residues, the Ramachandran angles [38], and the transform matrix between different local residue frames. Specifically, the local residue frame mentioned here represents the local coordinate system of an amino acid, characterizing the spatial position information of an amino acid. To build the local residue frame, AlphaFold2 [23] employs a Gram-Schmidt process on the backbone atom coordinates of a residue; and other methods, *e*.*g*., PiFold [14] and AlphaDesign [15], follow the method proposed by StructTrans [20] to set up a local frame for each residue. In this paper, we also adapt the Gram-Schmidt process, proposed by AlphaFold2 [23], to build a local residue frame for the vector field operator.

Based on the above hand-crafted features, previous methods made significant progress in protein inverse folding tasks. However, hand-crafted features were required by past methods because their networks could not extract geometric features. In this paper, the operators based on the transform matrix between local frames are proposed, allowing the network of VFN to extract geometric information, so hand-crafted features are no longer needed. Our method represents a new paradigm for extracting geometric information.

### 2.2 Inverse Protein Folding Problem

Engineered proteins have the opportunity to address many challenges in bio-science, biomedicine and chemistry, which attracted the interest of many researchers [4, 51, 2, 8, 41, 9]. The recent ML-based protein design paradigm [47] comprises two key steps. Firstly, a protein structure diffusion model is employed to generate a protein structure that fulfills the design specifications. Subsequently, the protein sequence is designed based on the generated protein structure. However, this latter step encounters a significant challenge referred to as the *inverse protein folding problem*:

Given a desired protein conformation, represented as the backbone atoms coordinates of each residue in a protein *χ* = {**x**_*i,k*_ ∈ ℝ^3^ : 1 ≤ *i* ≤ *N*, 1 ≤ *k* ≤ 4} (where *i* represents the *i*-th residue in the protein with *N* amino acids in total, and *k* denotes the *k*-th atoms in a residue backbone, 4 atoms in total), a method needs to predict the type of each amino acid *C* = {*c*_*i*_ ∈ ℝ} (*c*_*i*_ is the class index for residue, 20 types in total), which can be folded into the desired conformation. Previous machine-learning methods attempted to address this problem with two paradigms: The auto-regressive method [9] and the one-shot method [14]. In this work, the one-shot method is adopted for simplicity.

### 2.3 Protein Structure Modeling Methods

Based on the protein inverse folding problem, many methods have been proposed [43, 21, 28, 48, 36, 35, 10, 13, 6, 9, 20, 31, 22, 44, 18, 33, 46, 42, 3, 27, 17, 24, 4, 30, 32, 19, 11, 25, 26]. Those methods can be categorized into distance map-based [7], voxel-based [49, 37, 40] and GNN-based methods. Recently, GNN-based methods dominate the protein inverse folding problem; they are introduced as follows. For GNN-based methods, a protein is usually represented as a graph based on *k*-nearest neighbor on euclidean distance. The node and edge representations of the graph are initialized with hand-crafted geometric features. A message passing neural network (MPNN) [16] can then be applied to these graph representations to model the protein structure. GraphTrans [20] first proposed such paradigm and designed a transformer-based [45] GNN model. Then, GVP [22] proposed a novel geometric vector perception module, which also uses vector representation. Recently, more hand-crafted geometric feature-based methods have been proposed, *e*.*g*., AlphaDesign [21], ProteinMPNN [9], PiFold [14], and the performance has been continuously improved. Among them, PiFold is the current state-of-the-art method, partly because of the more comprehensive hand-crafted featurizer. However, as mentioned, VFN can extract the geometric feature through the vector field operator; such hand-crafted geometric feature is no longer needed.

## 3 Methodology

In this section, we present our protein structure modeling method, VFN. In VFN, as shown in Fig. 1 (A), a protein is represented as a *k*-NN graph [20]. Meanwhile, the implicit representations for nodes and edges are initialized. The resulting graph is fed into an *L*-layer message-passing neural network (MPNN) [16] to propagate messages and aggregate information for modeling the protein structure. The pipeline of each layer is the same. Within a layer, the node and edge representations are sequentially updated through three modules: the node aggregation module, the edge aggregation module, and the global context module. The *node aggregation module* aggregates information from adjacent nodes and edges to update node representations. The *edge aggregation module* aggregates the nodes incident to an edge to update the edge representation. The *global context module* updates each node by aggregating all node representations.

Particularly, unlike many previous methods [14, 9] that only utilize scalar hidden representations, we maintain a set of hidden geometric vector representations for each residue. In the node aggregation module, those vector representations are utilized to construct a vector field for extracting geometric features between residues. Moreover, SE(3)-invariance is naturally achieved in VFN, due to all the hidden geometric vectors being calculated under the local residue frame, not global frame, and there are not GVP-like [22] limitations (§2.3) on the network operation in VFN. Details are introduced as follows.

### 3.1 Protein Graph Initialization

In VFN, a protein is represented as a *k*-NN graph 𝒢 = (𝒮, ℰ). Specifically, every residue in the protein will be taken as a node, 𝒮 = {*s*_*i*_ : 1 ≤ *i* ≤ *N*}. Then, for each node, the edge is constructed between it and its neighbors, written as ℰ = {*e*_*i,j*_}_*j*∈Topk(*i*)_, where Topk(*i*) returns the index of *k*-nearest neighbor for *i*-th residue sorted on the euclidean distance. After constructing the protein graph, we compute five initial features, 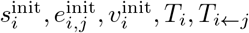, for the inputs and operations of VFN. Those features are introduced following.

Firstly, we follow the method proposed by PiFold [14] to initialize the *d*-dimensional scalar representation of node 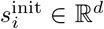. Specifically, a MLP layer is applied on the angle and distance between atoms inside the residue to generate 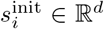. For the edge scalar representation 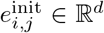, we train a group of statistic learnable weights as the 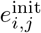. Notably, a complex geometric featurizer is required for initializing the 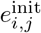 in many previous works [14, 21, 9]. Here, due to VFN can capture the geometric feature, the requirement of using a featurizer is eliminated. Subsequently, the *V* -channel vector representation 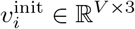 for each node consists of two components, which are initialized separately. The first part of 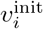 is initialized to the coordinates of backbone atoms **x**_*i*_ ∈ ℝ^4*×*3^, while the remaining part is initialized by a set of static learnable weights. The 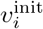 is represented under the *i*-th local residue frame *T*_*i*_, which is established by:

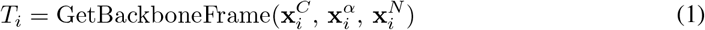

where *T*_*i*_ denotes the Euclidean transform from the *i*-th local residue frame to a global reference frame. GetBackboneFrame is a Gram–Schmidt process for setting up the local residue frame, illustrated in supplementary, proposed by AlphaFold2 [23].

With *T*_*i*_, we can calculate the transform matrix between different local residue frames for each edge in ℰ. Those can be written as:

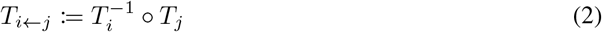

*T*_*i←j*_ represents the transform matrix for transforming a point coordinate from *j*-th local residue frame to *i*-th local residue frame; The ○ operation represents the composition of Euclidean transformations for two transform matrices; 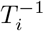 denotes the inverse transform of *T*_*i*_. Those transform matrices will be used to transform the hidden vector representation between different local residue frames later.

### 3.2 Geometric Vector Field Network

VFN is a *L*-layers sequential network, which takes 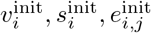 as the first layer input. All the layers in VFN are the same, and for a single layer in VFN, we denote the layer inputs as 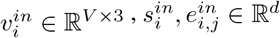 and the outputs as 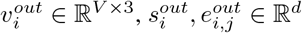.

#### Constructing the vector field

As shown in Fig.1(B) at the layer beginning, a vector field 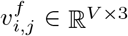 is constructed for each edge in ℰ to extract geometric feature for the node and edge aggregation module following. The way of building the vector field for an edge is transforming the neighbor hidden vector representation 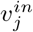 from the neighbor local frame *T*_*j*_ to its local frame *T*_*i*_ as points, and concatenating with 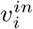 to generate an intermediate variable 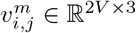. Then, the variable 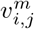 is sent into a vector linear layer V-Linear to generate the 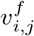, denoted as

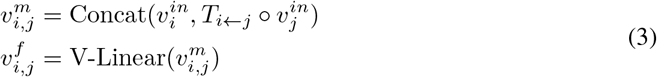

where the process of vector linear layer V-Linear is illustrated in supplementary material. After that, 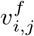 is transformed into a scalar representation *g*_*i,j*_ to extract geometric attribute for subsequent modules. This transformation 𝒯 decomposes 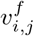 into the unit direction vectors and the vector lengths, which are encoded by a modified Gaussian radial basis function *φ*, similar to previous works [50]. The process of 𝒯 is written below:

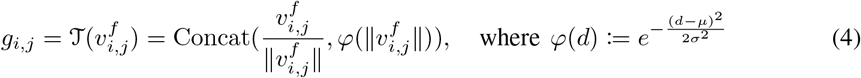

where *µ, σ* ∈ ℝ^*B*^ are sets of learnable paramters; *B* is the number of the Gaussian kernel; *g*_*i,j*_ ∈ ℝ^(3+*B*)*V*^ is the resulting geometric attribute.

#### Node aggregation module

After obtaining the geometric attribute *g*_*i,j*_ for each edge, the node aggregation module is employed to update the node representation 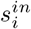 and 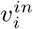 by synthesizing the information of 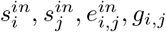 via a transformer-base [45, 14] mechanism. Firstly, those variables are sent into a MLP layer to generate the attention weights 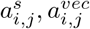 for scale attention and vector attention, respectively, written as:

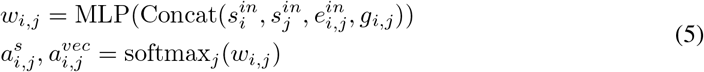

where 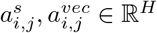 is the attention weights for scalar feature and vector field, respectively; *H* is the number of heads in the multi-head attention mechanism [45].

Next, we calculate the scalar value 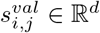 and the vector field value 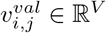, separately, for the multi-head attention. For the scale attention, the 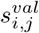 is firstly calculated by sending the 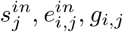 into a MLP layer and then summed weighted according to the 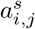. Similarly, for the vector attention, the 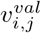 is generated by transforming the neighbour vector representation 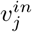 to centre local frame *T*_*i*_ and then applying another V-Linear layer on it; the resulting vector variable is summed weighted based on the 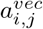. Those processes can be written as:

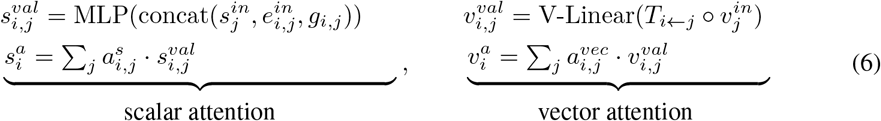

The 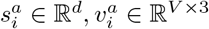 are the resulting scalar and vector features from the multi-head attention, respectively. Notably, there are *H* heads in the attention; the channel of the value 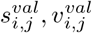 is split to *H* group and weighted summed with corresponding head in 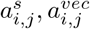, independently. Next, 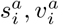 is sent to a MLP layer, independently:

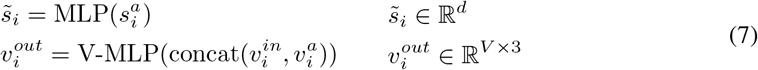

Specifically, here, we design a unique MLP layer for the vector field, termed V-MLP. In the V-MLP, there are two V-Linear layers and a direction activation function *η*, termed vector cosine-similarity activation, basing on the cosine similarity between the hidden vectors direction and the direction of a group statistic learnable vector, illustrated as:

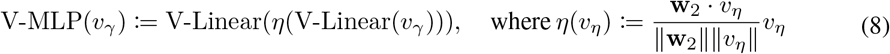

where 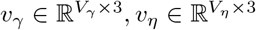 is the *V*_*γ*_-channel and *V*_*η*_-channel input of V-MLP and *η*, respectively, and 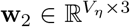 is a group of learnable vector weight. We refer readers to supplementary material for more detailed definitions of the V-MLP.

#### Edge aggregation module

Next, the 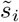 and *g*_*i,j*_ is used to update the edge representation 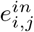 via a MLP layer, as shown in:

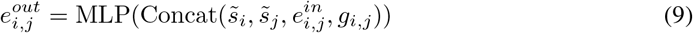

#### Global context module

Similar to PiFold [14] and GCA [44], a global context module is also employed to extract the global information for 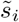, in which, the 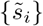 is averaged globally to a global representation *g* ∈ ℝ^*d*^. Then, *g* is used to update 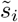 via a gating mechanism, defined as:

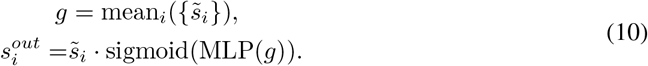

### 3.3 Decoder and Loss Function

Following PiFold, VFN predicts the type of residue in a one-shot style [14]. The decoder is extremely simple, consisting of only a single Linear layer on the final layer node representation *s*_*i*_. During training, the cross entropy loss is employed to supervise our model.

## 4 Experiments

In this section, we conduct extensive experiments on various datasets including the CATH 4.2 dataset [34] and the TS50 and TS500 datasets [28] to evaluate the performance of our proposed method. The results show that VFN achieves superior recovery scores compared to state-of-the-art methods, and demonstrates good scalability on a larger dataset.

### 4.1 Implementation Details

#### Datasets

Unless specified, our experiments are conducted on the CATH 4.2 [34] dataset using the same data splitting as previous works such as GVP [22] and PiFold [14]. The dataset consists of 18,024 proteins for training, 608 for validation, and 1120 for testing. During the evaluation, we also test our model on two smaller datasets, TS50 and TS500 [22, 14], to validate the generalizability. Furthermore, we also create another larger training set by incorporating data from the PDB [5]. We apply the same strategy as in [50] to collect and filter structures. Additionally, the proteins with sequences highly similar to test set proteins are also removed. By using the expanded dataset, we are able to scale up the VFN.

#### Training

All models are trained with batch size 8 and are optimized by AdamW with a weight decay of 0.1. We apply a OneCycle scheduler with a learning rate of 0.001 and train our model for a total of 100,000 iterations.

#### Model settings

Unless otherwise specified, the number of layers *L* is set to 10 and the number of vector *V* is set to 32. The dimension of scalar hidden representation *d* is 128.

### 4.2 Main results

#### Experimental results on CATH

In Table 1, we present the perplexity and recovery score results for VFN, compared to previous works such as [20, 22, 44, 18, 14], across various subsets of the CATH 4.2 dataset. Among the subsets considered in our analysis, the “Short” subset refers to proteins with a length of up to 100 amino acids, while the “Single chain” subset exclusively includes single chain proteins.

**Table 1:** Experimental results comparison on the CATH dataset. All baselines are reproduced by [14]. “Ours^+^” refers to a larger version of our proposed method that is trained on a larger dataset. “†” refers to the usage of a trick that assigns sequence labels to the residues with missing coordinates.

| Model | Venue | Perplexity↓ |  |  | Recovery(%) ↑ |  |  | CATH version |
| --- | --- | --- | --- | --- | --- | --- | --- | --- |
|  |  | Short | Single-chain | All | Short | Single-chain | All |  |
| StructGNN [20] | NeurIPS’19 | 8.29 | 8.74 | 6.40 | 29.44 | 28.26 | 35.91 | 4.2 |
| GraphTrans [20] | NeurIPS’19 | 8.39 | 8.83 | 6.63 | 28.14 | 28.46 | 35.82 | 4.2 |
| GCA [44] | arXiv’22 | 7.09 | 7.49 | 6.05 | 32.62 | 31.10 | 37.64 | 4.2 |
| GVP [22] | ICLR’20 | 7.23 | 7.84 | 5.36 | 30.60 | 28.95 | 39.47 | 4.2 |
| GVP-large | ICLR’20 | 7.68 | 6.12 | 6.17 | 32.60 | 39.40 | 39.20 | 4.3 |
| AlphaDesign [15] | arXiv’22 | 7.32 | 7.63 | 6.30 | 34.16 | 32.66 | 41.31 | 4.2 |
| ESM-IF [18] | ICML’22 | 8.18 | 6.33 | 6.44 | 31.30 | 38.50 | 38.30 | 4.3 |
| ProteinMPNN [9] | Science’22 | 6.21 | 6.68 | 4.61 | 36.35 | 34.43 | 45.96 | 4.2 |
| ProteinMPNN† [9] | Science’22 | - | - | 4.61 | - | - | 50.80 | 4.2 |
| PiFold[14] | ICLR’22 | 6.04 | 6.31 | 4.55 | 39.84 | 38.53 | 51.66 | 4.2 |
| Ours | this work | <u>5.55</u> | <u>5.55</u> | <u>4.14</u> | **42.94** | 40.83 | 54.74 | 4.2 |
| Ours<sup>+</sup> | this work | **5.31** | **5.32** | **3.66** | <u>42.76</u> | **41.77** | **57.14** | 4.2 |

Compared to the other methods, our proposed method shows superior recovery scores on all three datasets. To demonstrate the scalability of our model, we train a model on a larger dataset labeled as “Ours^+^”. The experiment results show that “Ours^+^” significantly outperforms “Ours”, highlighting the effectiveness of our approach in handling larger and more diverse protein structures. The model of “Ours^+^” will be released to aid in the designing of engineered proteins.

Furthermore, we select a challenging sample (PDB 2KRT) from the CATH test set and use AlphaFold2 to predict its structure based on the predicted sequence. We then visually compared the predicted structure with the experimentally determined native structure, as shown in Fig. 2. The visualizations demonstrate that, compared with PiFold, VFN achieves a higher accuracy on structure recovery, with only minor differences from the experimentally determined protein structures. In Fig. 3, we further compare the overall performance differences between our model and PiFold.

**Figure 2:**
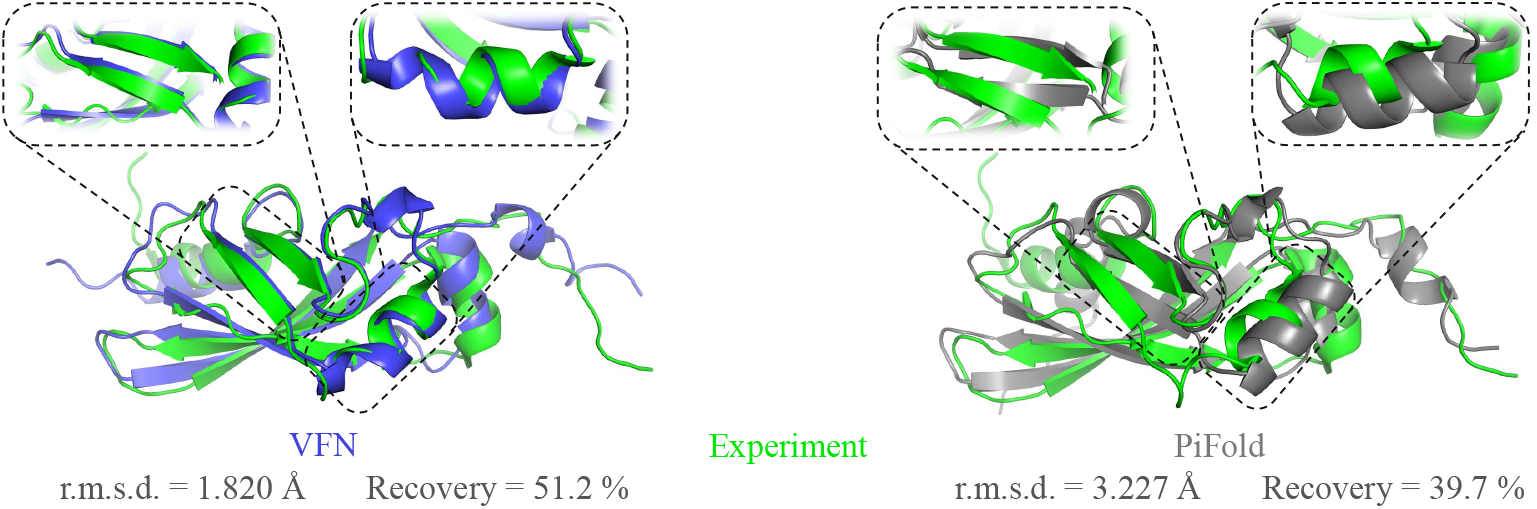
Visualization results of a challenging sample (PDB 2KRT). We use AlphaFold2 to recover the structure based on the predicted sequence and compare it against the experimentally determined ground-truth structure.

**Figure 3:**
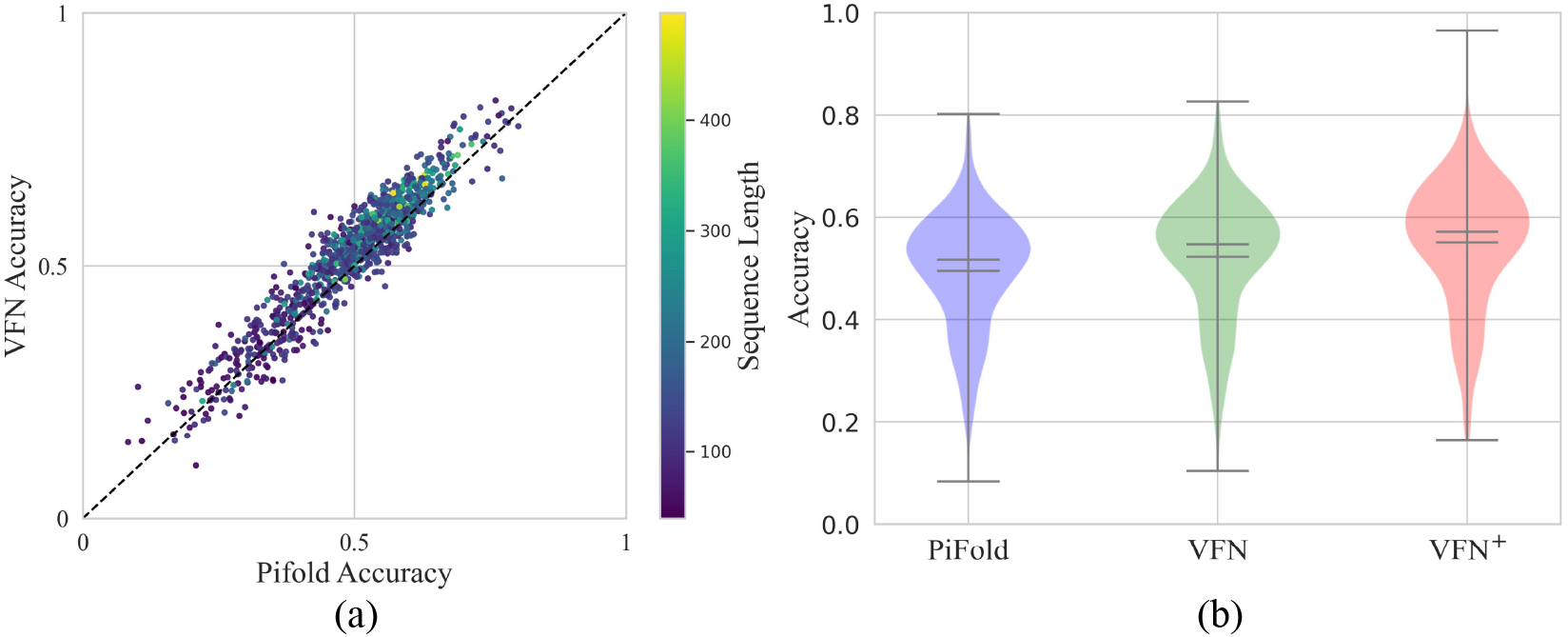
The performance comparison of VFN, VFN^+^ and PiFold. In Figure (a), each point represents a protein, its *x*- and *y*-coordinates represent the prediction accuracy of two models respectively, and its color corresponds to the sequence length. In Figure (b), we show the recovery accuracy distribution for different models on the CATH 4.2 test set [34].

#### Experimental results on TS50 and TS500

We further evaluate the performance of VFN on TS50 and TS500 datasets to examine the model’s generalizability. As shown in Table 2, we follow PiFold [14] and report the perplexity and median recovery on the TS50 and TS500 datasets. Results show that our method outperforms other state-of-the-art methods in terms of recovery scores, with a median recovery score of 63.74% on TS500 and 60.38% on TS50. Furthermore, our method achieves a lower perplexity than PiFold on both datasets, indicating better generalization ability.

**Table 2:**
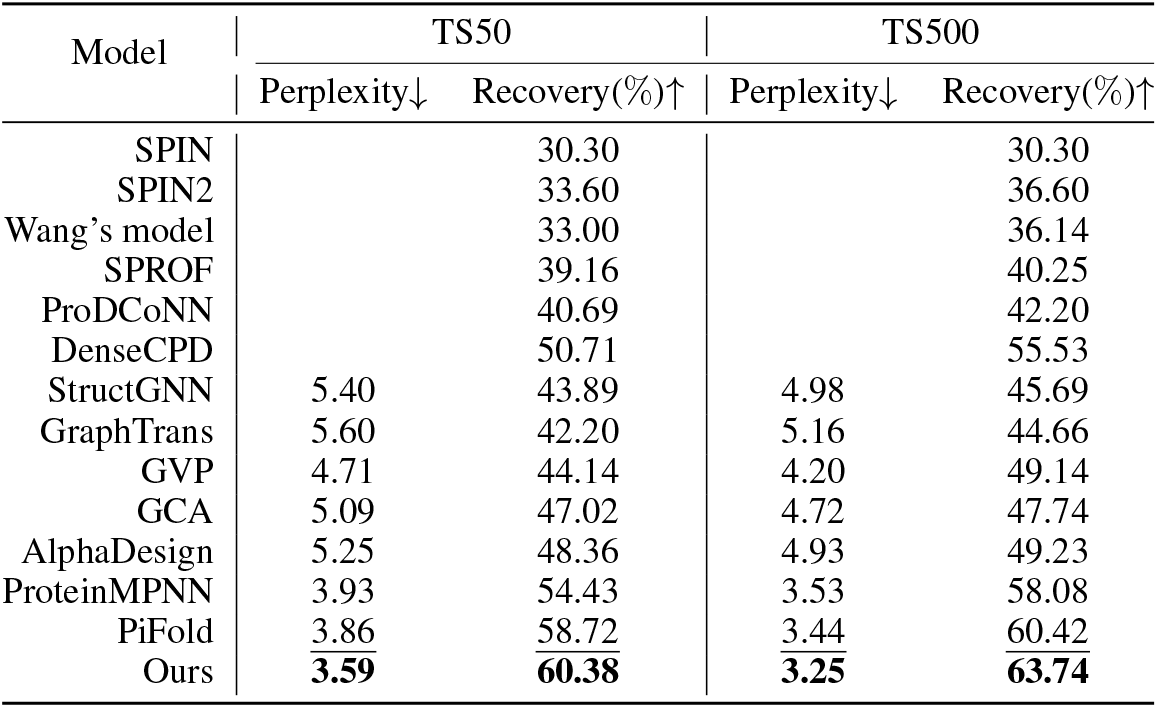
Experimental results on TS50 and TS500.The optimal results are labeled in bold, and suboptimal results are underlined.

### 4.3 Ablations

In this section, we carefully investigate the design choices of vector field modules proposed here.

#### The number of layers

We investigate the impact of modifying the number of layers on recovery and perplexity in Table 3a. Increasing the number of layers from 5 to 15 results in a marginal improvement in recovery scores, with the highest recovery achieved at 54.72% for 12 layers. Ablation experiments without the edge featurizer show that with or without edge features, the performance is comparable. Especially when the number of layers reaches 15, it even achieves better results, indirectly proving the effectiveness of VFN and its potential for reducing reliance on hand-crafted features.

**Table 3:**
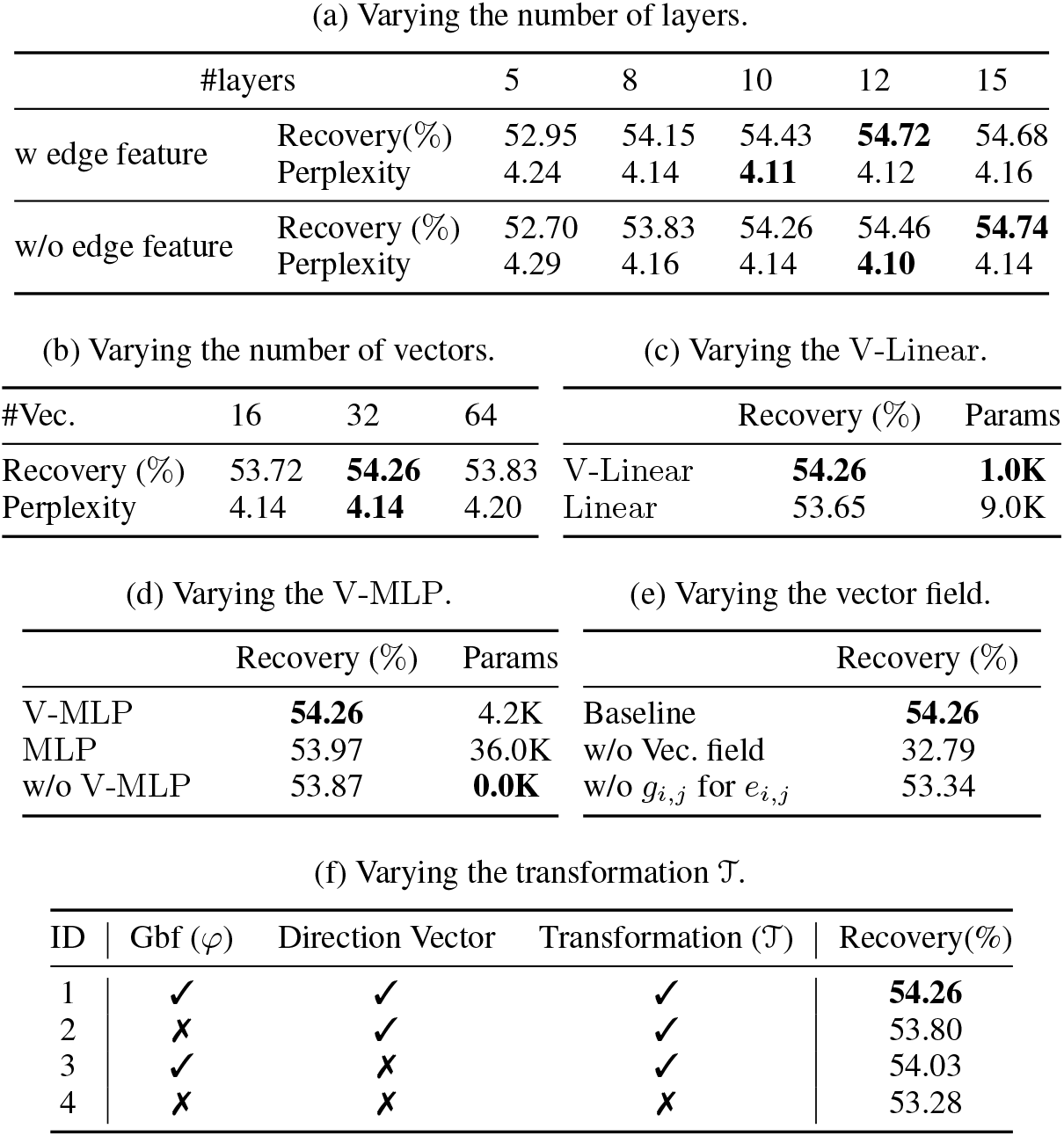
Ablation studies on the CATH 4.2 dataset. We use the default model settings unless otherwise specified. When calculating the number of parameters, we only count the number of parameters occupied by this module in one layer.

#### The number of vectors

The ablation study in Table 3b reveals that the most favorable outcome is obtained with a vector number of 32.

#### V-Linear

Table 3c demonstrates that replacing the original Linear with V-Linear reduces the number of parameters and improves model performance.

#### V-MLP

In Table 3d, we observe that using V-MLP outperforms using a regular MLP or not using it at all. Compared to using a regular MLP, V-MLP significantly reduces the parameter count.

#### Vector field design

Results in Table 3e show that removing the whole vector field module leads to a significant drop in recovery, indicating its importance in capturing protein folding patterns. The incorporation of the *g*_*i,j*_ in the edge aggregation module also has a substantial effect on the performance of the model.

#### Transformation 𝒯 design

We validate the effectiveness of Gaussian radial basis function *φ* and Direction Vector in the Transformation 𝒯, as shown in Table 3f. Furthermore, it illustrates that completely excluding the Transformation 𝒯 and directly flattening the vectors into scalar representations leads to an obvious performance decrease.

## 5 Conclusion

In this paper, we propose VFN, which effectively learns to extract geometric features, leading to state-of-the-art performance in inverse folding tasks. We demonstrate its ability to capture geometric features and minimize the need for complex manual feature engineering. This advancement offers a new paradigm for future geometric network designs.

## References

[1] Abien Fred Agarap. Deep learning using rectified linear units (ReLU). arXiv preprint arXiv:1803.08375, 2018.

[2] Rebecca F. Alford, Andrew Leaver-Fay, Jeliazko R. Jeliazkov Matthew J. O’Meara, Frank P. DiMaio, Hahnbeom Park, Maxim V. Shapovalov, P. Douglas Renfrew, Vikram K. Mulligan, and et al. Kappel, Kalli. The Rosetta all-atom energy function for macromolecular modeling and design. Computational and Structural Biotechnology Journal, 13(6):3031–3048, 2017.

[3] Namrata Anand, Raphael Eguchi, Irimpan I. Mathews, Carla P. Perez, Alexander Derry, Russ B. Altman, and Po-Ssu Huang. Protein sequence design with a learned potential. Nature Communications, 13(1):746, 2022.

[4] Ivan Anishchenko, Samuel J. Pellock, Tamuka M. Chidyausiku, Theresa A. Ramelot, Sergey Ovchinnikov, Jingzhou Hao, Khushboo Bafna, Christoffer Norn, Alex Kang, and et al. Bera, Asim K. De novo protein design by deep network hallucination. Nature, 600(7889):547–552, 2021.

[5] Stephen K. Burley, Helen M. Berman, Gerard J. Kleywegt, John L. Markley, Haruki Nakamura, and Sameer Velankar. Protein data bank (pdb): the single global macromolecular structure archive. Protein Crystallography: Methods and Protocols, pages 627–641, 2017.

[6] Yue Cao, Payel Das, Vijil Chenthamarakshan, Pin-Yu Chen, Igor Melnyk, and Yang Shen. Fold2seq: A joint sequence (1d)-fold (3d) embedding-based generative model for protein design. In International Conference on Machine Learning, pages 1261–1271. PMLR, 2021.

[7] Sheng Chen, Zhe Sun, Lihua Lin, Zifeng Liu, Xun Liu, Yutian Chong, Yutong Lu, Huiying Zhao, and Yuedong Yang. To improve protein sequence profile prediction through image captioning on pairwise residue distance map. Journal of Chemical Information and Modeling, 60(1):391–399, 2019.

[8] Bassil I. Dahiyat and Stephen L. Mayo. Probing the role of packing specificity in protein design. Proceedings of the National Academy of Sciences, 94(19):10172–10177, 1997.

[9] Justas Dauparas, Ivan Anishchenko, Nathaniel Bennett, Hua Bai, Robert J. Ragotte, Lukas F. Milles, Basile IM Wicky, Alexis Courbet, Rob J. de Haas, and et al. Bethel, Neville. Robust deep learning–based protein sequence design using proteinmpnn. Science, 378(6615):49–56, 2022.

[10] Wenze Ding, Kenta Nakai, and Haipeng Gong. Protein design via deep learning. Briefings in Bioinformatics, 23(3), 2022.

[11] Baldwin Dumortier, Antoine Liutkus, Clément Carré, and Gabriel Krouk. Petribert: Augmenting bert with tridimensional encoding for inverse protein folding and design. BioRxiv, pages 2022–08, 2022.

[12] Richard Evans Michael O’Neill, Alexander Pritzel, Natasha Antropova, Andrew Senior, Tim Green, Augustin Žídek, Russ Bates, Sam Blackwell, Jason Yim, Olaf Ronneberger, Sebastian Bodenstein, Michal Zielinski, Alex Bridgland, Anna Potapenko, Andrew Cowie, Kathryn Tunyasuvunakool, Rishub Jain, Ellen Clancy, Pushmeet Kohli, John Jumper, and Demis Hassabis. Protein complex prediction with alphafold-multimer. BioRxiv, 2021.

[13] Wenhao Gao, Sai Pooja Mahajan, Jeremias Sulam, and Jeffrey J. Gray. Deep learning in protein structural modeling and design. Patterns, 1(9):100142, 2020.

[14] Zhangyang Gao, Cheng Tan, and Stan Li.PiFold: Toward effective and efficient protein inverse folding. International Conference on Learning Representations, 2022.

[15] Zhangyang Gao, Cheng Tan, and Stan Z Li. Alphadesign: A graph protein design method and benchmark on alphafolddb. arXiv preprint arXiv:2202.01079, 2022.

[16] Justin Gilmer, Samuel S. Schoenholz, Patrick F. Riley, Oriol Vinyals, and George E. Dahl.Message passing neural networks. Machine Learning Meets Quantum Physics,pages 199–214, 2020.

[17] Joe G. Greener, Lewis Moffat, and David T. Jones. Design of metalloproteins and novel protein folds using variational autoencoders. Scientific Reports, 8(1):16189, 2018.

[18] Chloe Hsu, Robert Verkuil, Jason Liu, Zeming Lin, Brian Hie, Tom Sercu, Adam Lerer, and Alexander Rives. Learning inverse folding from millions of predicted structures. In International Conference on Machine Learning, pages 8946–8970. PMLR, 2022.

[19] Bin Huang, Tingwen Fan, Kaiyue Wang, Haicang Zhang, Chungong Yu, Shuyu Nie, Yangshuo Qi, Wei-Mou Zheng, Jian Han, and et al. Fan, Zheng. Accurate and efficient protein sequence design through learning concise local environment of residues. Bioinformatics, page btad122, 2023.

[20] John Ingraham, Vikas Garg, Regina Barzilay, and Tommi Jaakkola. Generative models for graph-based protein design. Advances in Neural Information Processing Systems, 32, 2019.

[21] Michael Jendrusch, Jan Korbel, and Kashif Sadiq. Alphadesign: A de novo protein design framework based on AlphaFold. BioRxiv, pages 2021–10, 2021.

[22] Bowen Jing, Stephan Eismann, Patricia Suriana, Raphael Townshend, and Ron Dror. Learning from protein structure with geometric vector perceptrons. International Conference on Learning Representations, 2020.

[23] John Jumper, Richard Evans, Alexander Pritzel, Tim Green, Michael Figurnov, Olaf Ronneberger, Kathryn Tunyasuvunakool, Russ Bates, Augustin Žídek, and et al. Potapenko, Anna. Highly accurate protein structure prediction with alphafold. Nature, 596(7873):583–589, 2021.

[24] Mostafa Karimi, Shaowen Zhu, Yue Cao, and Yang Shen. De novo protein design for novel folds using guided conditional wasserstein generative adversarial networks. Journal of Chemical Information and Modeling, 60(12):5667–5681, 2020.

[25] Alex J. Li, Mindren Lu, Israel Desta, Vikram Sundar, Gevorg Grigoryan, and Amy E. Keating. Neural network-derived potts models for structure-based protein design using backbone atomic coordinates and tertiary motifs. Protein Science, 32(2):e4554, 2023.

[26] Alex J. Li, Vikram Sundar, Gevorg Grigoryan, and Amy E. Keating. Terminator: a neural framework for structure-based protein design using tertiary repeating motifs. arXiv preprint arXiv:2204.13048, 2022.

[27] Jie Li and Patrice Koehl. 3d representations of amino acids—applications to protein sequence comparison and classification. Computational and Structural Biotechnology Journal, 11(18):47–58, 2014.

[28] Zhixiu Li, Yuedong Yang, Eshel Faraggi, Jian Zhan, and Yaoqi Zhou. Direct prediction of profiles of sequences compatible with a protein structure by neural networks with fragment-based local and energy-based nonlocal profiles. Proteins: Structure, Function, and Bioinformatics, 82(10):2565–2573, 2014.

[29] Zeming Lin, Halil Akin, Roshan Rao, Brian Hie, Zhongkai Zhu, Wenting Lu, Nikita Smetanin, Allan dos Santos Costa, Maryam Fazel-Zarandi, Tom Sercu, and et al. Candido, Sal. Language models of protein sequences at the scale of evolution enable accurate structure prediction. BioRxiv, 2022.

[30] Yufeng Liu, Lu Zhang, Weilun Wang, Min Zhu, Chenchen Wang, Fudong Li, Jiahai Zhang, Houqiang Li, Quan Chen, and Haiyan Liu. Rotamer-free protein sequence design based on deep learning and self-consistency. Nature Computational Science, 2(7):451–462, 2022.

[31] Jack B. Maguire, Daniele Grattarola, Vikram Khipple Mulligan, Eugene Klyshko, and Hans Melo. Xenet: Using a new graph convolution to accelerate the timeline for protein design on quantum computers. PLoS Computational Biology, 17(9):e1009037, 2021.

[32] Matt McPartlon, Ben Lai, and Jinbo Xu. A deep SE(3)-equivariant model for learning inverse protein folding. BioRxiv, pages 2022–04, 2022.

[33] James O’Connell, Zhixiu Li, Jack Hanson, Rhys Heffernan, James Lyons, Kuldip Paliwal, Abdollah Dehzangi, Yuedong Yang, and Yaoqi Zhou. Spin2: Predicting sequence profiles from protein structures using deep neural networks. Proteins: Structure, Function, and Bioinformatics, 86(6):629–633, 2018.

[34] Christine Orengo, Alex Michie, Susan Jones, David Jones, Mark Swindells, and Janet Thornton. Cath–a hierarchic classification of protein domain structures. Structure, 5(8):1093–1109, 1997.

[35] Sergey Ovchinnikov and Po-Ssu Huang. Structure-based protein design with deep learning. Current Opinion in Structural Biology, 65:136–144, 2021.

[36] Robin Pearce and Yang Zhang. Deep learning techniques have significantly impacted protein structure prediction and protein design. Current Opinion in Structural Biology, 68:194–207, 2021.

[37] Yifei Qi and John ZH Zhang. Densecpd: improving the accuracy of neural-network-based computational protein sequence design with densenet. Journal of Chemical Information and Modeling, 60(3):1245–1252, 2020.

[38] GN. T. Ramachandran and V. Sasisekharan. Conformation of polypeptides and proteins. Advances in Protein Chemistry, 23:283–437, 1968.

[39] Carol A. Rohl, Charlie EM. Strauss, Kira MS. Misura, and David Baker. Protein structure prediction using rosetta.In Methods in enzymology, volume 383, pages 66–93. Elsevier, 2004.

[40] Raghav Shroff, Austin Cole, Barrett Morrow, Daniel Diaz, Isaac Donnell, Jimmy Gollihar, Andrew Ellington, and Ross Thyer. A structure-based deep learning framework for protein engineering. BioRxiv, 2019.

[41] Arthur G. Street and Stephen L. Mayo. Computational protein design. Structure, 7(5):R105–R109, 1999.

[42] Alexey Strokach, David Becerra, Carles Corbi-Verge, Albert Perez-Riba, and Philip M. Kim. Ftast and flexible protein design using deep graph neural networks. Cell Systems, 11(4):402–411, 2020.

[43] Alexey Strokach and Philip M. Kim. Deep generative modeling for protein design. Current Opinion in Structural Biology, 72:226–236, 2022.

[44] Cheng Tan, Zhangyang Gao, Jun Xia, and Stan Li. Generative de novo protein design with global context. arXiv preprint arXiv:2204.10673, 2022.

[45] Ashish Vaswani, Noam Shazeer, Niki Parmar, Jakob Uszkoreit, Llion Jones, Aidan N. Gomez, Łukasz Kaiser, and Illia Polosukhin. Attention is all you need. Advances in Neural Information Processing Systems, 30, 2017.

[46] Jingxue Wang, Huali Cao, John ZH Zhang, and Yifei Qi. Computational protein design with deep learning neural networks. Scientific Reports, 8(1):1–9, 2018.

[47] Joseph L Watson, David Juergens, Nathaniel R Bennett, Brian L Trippe, Jason Yim, Helen E Eisenach, Woody Ahern, Andrew J Borst, Robert J Ragotte, Lukas F Milles, et al. Broadly applicable and accurate protein design by integrating structure prediction networks and diffusion generative models. bioRxiv, pages 2022–12, 2022.

[48] Zachary Wu, Kadina E. Johnston, Frances H. Arnold, and Kevin K. Yang. Protein sequence design with deep generative models. Current Opinion in Structural Biology, 65:18–27, 2021.

[49] Yuan Zhang, Yang Chen, Chenran Wang, Chun-Chao Lo, Xiuwen Liu, Wei Wu, and Jinfeng Zhang. Prodconn: Protein design using a convolutional neural network. Proteins: Structure, Function, and Bioinformatics, 88(7):819–829, 2020.

[50] Gengmo Zhou, Zhifeng Gao, Qiankun Ding, Hang Zheng, Hongteng Xu, Zhewei Wei, Linfeng Zhang, and Guolin Ke. Uni-mol: A universal 3d molecular representation learning framework. 2023.

[51] Jianfu Zhou, Alexandra E. Panaitiu, and Gevorg Grigoryan. A general-purpose protein design framework based on mining sequence–structure relationships in known protein structures. Proceedings of the National Academy of Sciences, 117(2):1059–1068, 2020.

